# TabVI: Leveraging Lightweight Transformer Architectures to Learn Biologically Meaningful Cellular Representations

**DOI:** 10.1101/2025.02.13.637984

**Authors:** Aditi Chandrashekar, Rohan Gala, Andreas Tjärnberg, Saniya Khullar, Grace Huynh, Mariano Gabitto

## Abstract

Transformer-based foundation models are changing the landscape of natural language processing (NLP), computer vision, and audio, achieving human-level performance across a variety of tasks. Extending these models to single-cell genomics holds significant potential for revealing the cellular and molecular perturbations associated with disease. However, unlike the sequential structure of language, the functional organization of genes is hierarchical and modular. This fundamental difference necessitates the development of meaningful feature selection strategies to adapt NLP transformer architectures effectively. In contrast to many large-scale foundation models, probabilistic models have shown success in learning complex cellular representations from single-cell datasets. In this work, we present TabVI, a probabilistic deep generative model that leverages tabular transformer architectures to improve latent embedding learning. We validate TabVI’s performance in cell type annotation and integration benchmarks. We demonstrate that TabVI improves performance across down-stream tasks and is robust to scaling dataset sizes, producing interpretable, sample-specific feature attention masks. TabVI is a lightweight, scientifically-meaningful, transformer architecture for single-cell analysis that excels where large scale foundation models are less effective.

## 1 Introduction

Large-scale foundation models (FM) Bommasani et al. (2021) pre-trained on vast corpus of data are revolutionizing natural language processing, reaching human-level performance across wide spectrum of applications Devlin et al. (2019); OpenAI et al. (2024); Dubey et al. (2024); Vyas et al. (2023); Gardner et al. (2024); Oquab et al. (2024); Dosovitskiy et al. (2021); Wang et al. (2024). FMs have shown impressive generalization capabilities not only when fine-tuned for downstream tasks, but also when applied in zero-shot paradigms, suggesting the models have captured the underlying structure of language Yin et al. (2019); Brown et al. (2020); Mercea et al. (2022); Li et al. (2024); Chen et al. (2024). Attention-based transformer architectures are at the core of these models Vaswani et al. (2017). Using such models as a foundation upon which other models can be built is a promising endeavor for scientific machine learning, in which domain-specific scientific problems are solved by using tools from machine learning Subramanian et al. (2023); Ho (2024); H et al. (2023); Song et al. (2023). However, it is not known how broadly this methodological approach can be applied or how to readily interpret parameters within transformer architectures or their intermediate layers.

While transformer-based architectures and pre-training strategies have been successful for vision and language tasks, the creation and application of foundation models to single cell genomics datasets remains challenging H et al. (2024); Zheng Y (2023); Y et al. (2023); Alsabbagh et al. (2023); Boiarsky et al. (2023); Kedzierska et al. (2023). The absence of a clear, sequential ordering of input data (genes), and the noisy and high dimensional nature of -omics data are key challenges in the creation of a single-cell FM. More precisely, NLP transformer architectures incorporate the position of each word within the input to generate context-aware positional embeddings for data processing. While successful when adapted for use in genomics H et al. (2024), genes lack a predefined ordering, rendering this approach less aligned with biological principles.

In contrast, variational autoencoder (VAE)Kingma & Welling (2022) -based probabilistic approaches focus on learning unsupervised representations and have been successful in the analysis of large-scale uni- and multi-modal -omics datasets Lopez et al. (2018); Ashuach* et al. (2023); A. et al. (2022). These models offer a principled approach to describe the data generating process, quantify the uncertainty of estimated quantities, and provide an interpretable description of model parameters, accounting for precise technical and biological factors. VAE based methods are widely used to analyze, annotate, and visualize single-cell RNA-seq (scRNA-seq) data due to their ability to handle high-dimensional, large scale data Lopez et al. (2018); Ashuach* et al. (2023); A. et al. (2022).VAEs have enabled the construction of a detailed and cohesive understanding of cellular structure within tissues in health and disease Gabitto et al. (2023). To build such cellular atlases, the heterogeneous patterns of gene expression need to be parsed and cataloged Tjärnberg et al. (2024), a challenging task given that genes act combinatorially, with a modular and hierarchical interdependence Barabási et al. (2011), a product of the common developmental origin of cells.

In this work, we bridge the gap between the probabilistic and foundation modeling paradigms by developing an approach that incorporates a lightweight transformer module within a VAE representation learning framework for the analysis of scRNA-seq. We harness a recently developed tabular transformer architecture that processes the input data by extracting complex relationships across input features, akin to the one needed in cellular biology given the modular organization of genes, and creating per-sample attention masks Arik & Pfister (2020). Next, we modify the internal layers of the transformer architecture to enable better sample-efficient performance. We adapt this architecture to work as an encoder network within a probabilistic graphical model, creating a tabular-transformer variational autoencoder. We show that the proposed approach improves representations learned by a commonly-used baseline probabilistic deep generative model A. et al. (2022) and an NLP-based transformer architecture H et al. (2024), and can be learnt in a sample efficient manner, while offering an interpretable view of features and the sample space. These results suggest a path forward for developing scientific foundation models in genomics that leverage biologically-inspired transformer-based architectures to enhance the interpretability of both representations and input features. We showcase the model’s abilities by considering a recent single-nucleus RNA-seq (snRNA-seq) data set from the Seattle Alzheimer’s disease brain Cell Atlas (SEA-AD) Gabitto et al. (2023), in which individual cells were profiled from heterogeneous brain donors that spanned the entire spectrum of Alzheimer’s disease (AD) states.

## 2 TabVI

TabVI builds upon the probabilistic framework of scVI Lopez et al. (2018) for representation learning. By incorporating both discrete and continuous latent variables, akin to the approach used in scAnVI Xu et al. (2019), we have developed a model we call TabAnVI. TabAnVI infers a latent cellular space and facilitates the prediction of cell type annotations. In this section, we outline the probabilistic model of TabVI alongside our encoder transformer architecture (Figure 1). We will briefly introduce TabAnVI at the end of this section, details about the model can be found in Appendix A.1.

**Figure 1.**
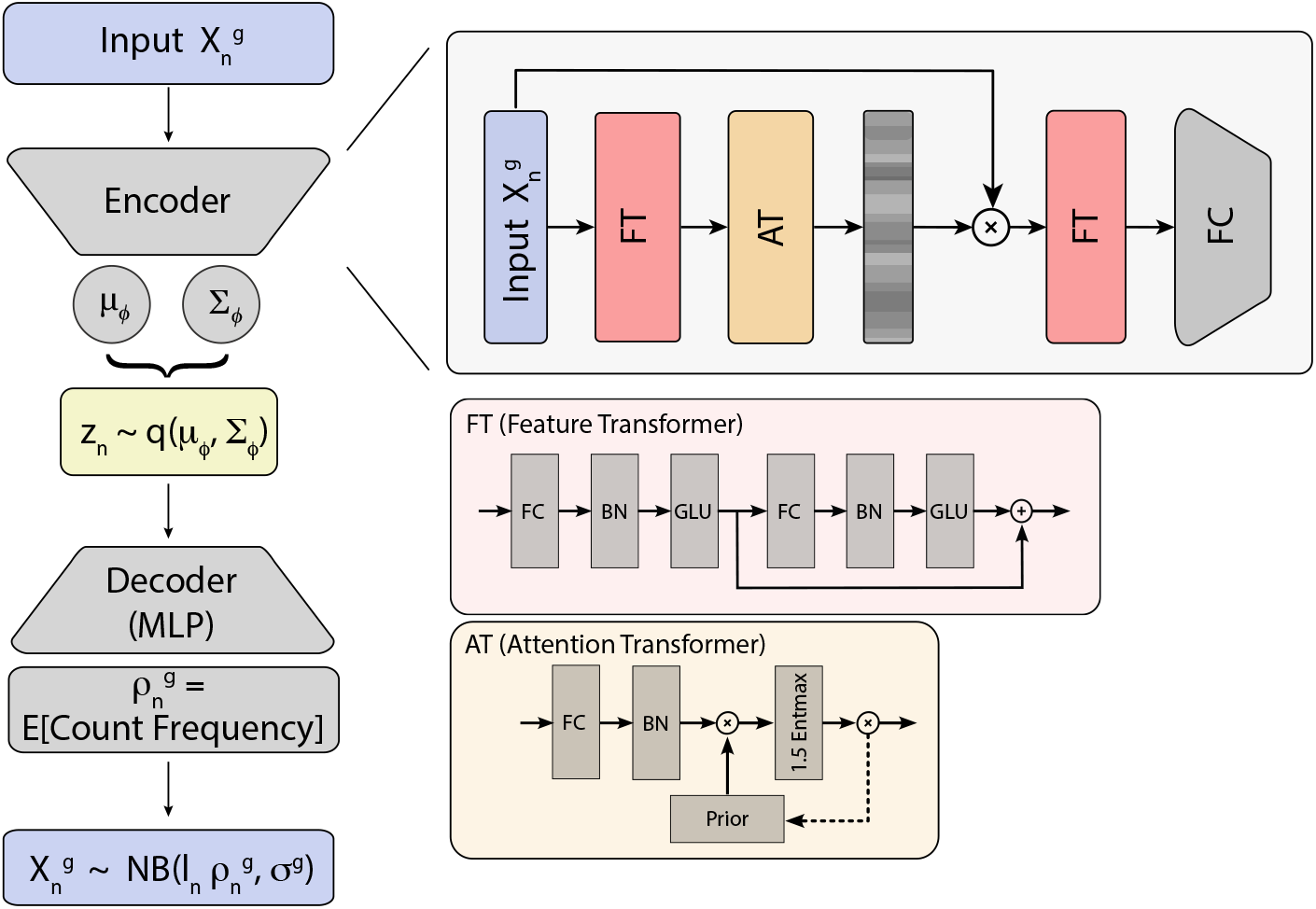
TabVI model architecture combines the latent space interpretability of VAEs with a sample efficient tabular feature transformer. Input gene expression values pass through feature and attentive transformer layers that provide masks on a per-sample basis. Masked features are encoded by a feature transformer to obtain each cell’s latent low dimensional representation. The representation is decoded through MLP layers to learn parameters of the probabilistic model with a reconstruction objective.

### 2.1 Probabilistic Model

The result of a single cell/nucleus RNA-sequencing (sc/nRNA-seq) experiment generates input data, 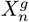, depicting the counts of mRNA molecules for the expression of gene *g* in cell *n*. Covariates associated with each cell (such as the batch that the cell was collected in) are denoted as *s*_*n*_. We assume that this experiment is generated under observational noise characterized by a zero-inflated negative binomial distribution with parameters *θ*_*g*_, *l*_*n*_, and 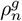, denoting gene specific dispersion, cell specific library size, and a mean expression in each cell per gene respectively. This noise model is selected to account for count overdispersion and the inherent sparsity of single cell measurements. Let *z*_*n*_ ∈ ℛ ^*d*^, with *d* ≪ *G* (with *G* being the total number of genes), represent a per-cell variable capturing the underlying cell biological state. The mean expression 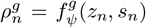 is calculated by a nonlinear transformation *f*_*ψ*_ : ℛ^d^ → ℛ^*G*^, with parameters *ψ*, from the latent cell state. A Bayesian prior is assumed for *z*_*n*_ ∈ ℛ^*K*^, with *K* as the dimension of the latent cell space, as a standard normal distribution. This entire model is summarized in equation 1.

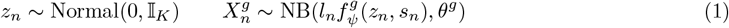

To perform inference on our model parameters and due to the intractable nature of their posterior, we resort to variational inference Blei et al. (2017). In particular, due to the nonlinearities present in the generative model, we use the variational autoencoder framework Kingma & Welling (2022) to perform inference. We assume a variational proposal of the form:

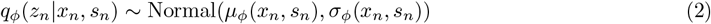

To learn the parameters of the model, we optimize the evidence lower bound (ELBO) as a function of the parameters *ψ* and *ϕ* using stochastic optimization Kingma & Ba (2017). In addition, *l*_*c*_ and *θ*_*g*_ are learnt as point estimates. The functions *f*_*ψ*_ and *g*_*ϕ*_ are commonly known as decoder and encoder respectively. In our baseline models, scVI and scAnVI models encoders are commonly parameterized by using multi-layer fully connected neural networks.

### 2.2 Transformer Architecture

To improve upon the baseline model and increase the expressiveness of the latent embedding while maintaining interpretability, we adopt a tabular transformer architecture Arik & Pfister (2020) as our encoder. This architecture comprises two primary components: a feature transformer and an attention transformer. The feature transformer maps the input *x* ∈ ℛ ^*B×G*^ (where *B* denotes batch size) to an embedding *Y*∈ ℛ ^*B×*(*D*+*A*)^, which is then split into two parts, *Y*_*D*_ and *Y*_*A*_, with dimensions *B× D* and *B × A*, respectively. *Y*_*D*_ is used to transform the input, while *Y*_*A*_ serves as the input for the attention transformer. The attention transformer maps its input *Y*_*A*_ ∈ ℛ^*B×A*^ to ℛ^*B×G*^, producing an instance-specific attention mask of the same size as the input data, which is used to re-weight the input. The re-weighted data is then processed by a feature transformer layer. Finally, the output of this layer is passed through a fully connected layer, yielding the posterior mean and variance parameters required for the VAE architecture, as depicted in Figure 1. Details on the architecture’s building blocks and parameter sizes are provided in Appendix A.2.

TabAnVI model is a tabular-transformer extension of the semi-supervised probabilistic algorithm scAnVI. scAnVI provides a principled way to create cell label annotations by enriching the latent structure of scVI with a per-cell categorical variable *c*_*n*_, representing the cell type. Then, each cell has a latent random variable *u*_*n*_, describing biological variability. These two variables are nonlinearly combined into *z*_*n*_, describing a cell type aware state of the cell.

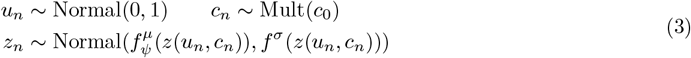

Similarly to TabVI and scVI, we use a variational autoencoder framework. Of note, after training, the variational proposals for the categorical variable can be used as a classifier as the latent cell type state can be derived directly from the model *q*_*ϕ*_(*c*_*n*_ |*z*_*n*_) = Cat(*g*_*ϕ*_(*z*_*n*_)), with *g* being a nonlinear function of the latent space with parameters *ϕ* (for a full description see Appendix A.1). For an ablation study on the components of this architecture, see Appendix A.6.

## 3 Related Work

## 4 Empirical Studies

### 4.1 Cell Type Prediction and Data Integration

TabVI aims to learn an interpretable low-dimensional latent space that captures biological variability and corrects for batch and technical effects present in the data. TabAnVI inherits this representation and learns a latent space containing continuous and discrete latent states that can be used for cell type annotation. To evaluate these capabilities, we considered a snRNA-seq data set profiling the human MTG in brain donors with AD from the Seattle Alzheimer’s Disease Brain Cell Atlas (section A.3). Brain donor information can be used as a batch covariate, *s*_*n*_, and condition the model on this information by providing this information for each cell. We randomly selected 80% of the SEA-AD dataset and subset it to the top 4k variable genes as input. We train TabVI, and use this trained model as a pre-trained architecture to train TabAnVI (more on the training procedure in appendix A.4), resulting in a latent representation that can be visualized through UMAP embedding McInnes et al. (2020) (Figure 2A).

**Figure 2.**
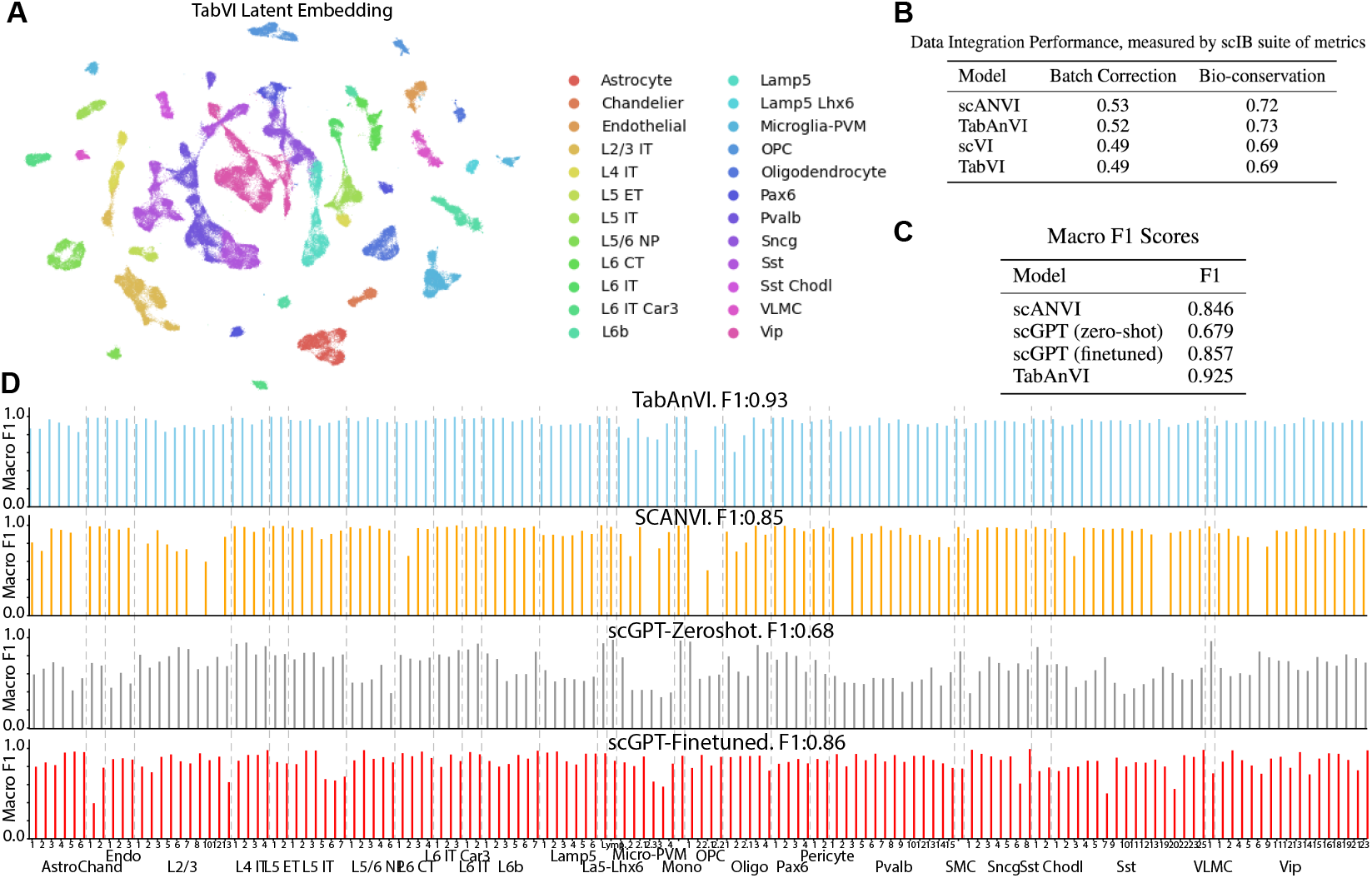
Benchmarking TabVI model across integration and annotation tasks. **A** TabVI’s latent representations of human middle temporal gyrus cells originating from SEA-AD donors spanning the entire spectrum of AD. Cells are color-coded by their subclass, where each subclass contains multiple cell types. **B, C** TabVI representations enable higher overall classification performance compared to scAnVI or scGPT representations, while maintaining scVI’s representation capabilities (as evaluated by using commonly used batch-integration metrics with donors treated as batches). **D** TabVI achieves high classification performance across cell types.

To evaluate latent space properties, we use the scIB platform Luecken et al. (2022), which provides metrics for assessing dataset integration by measuring batch correction (batch-correction, in our case multiple donors) and the preservation of biological variability (bio-conservation, in our case cell type label). In this evaluation, we examine whether incorporating our transformer architecture affects dataset integration compared to the baseline models, scVI and scAnVI. We find that our enhanced models perform comparably to the parent models (Figure 2B). When cell type labels are incorporated into the models (scAnVI and TabAnVI), batch-correction performance remains consistent, while bio-conservation improves significantly, a natural result of training both models with ground truth biological information.

We assess the models’ performance in predicting cell types when trained on 4000 genes, using F1 macro annotation scores to measure their ability to classify cell types in the remaining 20% of the dataset, which the models were not trained on. In this extended comparison, we evaluated TabAnVI against scAnVI, scGPT (in zero-shot mode), and scGPT (fine-tuned to our cell type labels) for cell type annotation (Figure 2C). TabAnVI outperformed the other models, achieving an F1 score of 0.925, while the next best, scGPT (finetuned), reached 0.857. Notably, both versions of scGPT classified all cell types with varying success (red and gray bars), whereas scVI failed to classify several cell types. TabAnVI showed strong performance across most cell types, missing only one OPC subtype.

These computational experiments underscore TabVI’s ability to learn a meaningful latent representation of single-nucleus data and accurately annotate granular cell types, outperforming even a state-of-the-art foundation model.

### 4.2 Sample and Feature Scaling Properties

FM architectures are composed of multiple transformer layers, imposing a high computational burden and requiring vast datasets for training. To study data- and feature-scaling capabilities of our architecture, we build upon our previous computational experiments (section 4.1) and evaluate TabVI’s annotation performance by varying the number of input features (genes) and dataset size. We compare TabVI performance against scAnVI by scaling the number of training samples, *N* (with *N* ∈ {50%, 75%, 100%} of the total dataset), and number of genes, *G*, (with *G* ∈ {1000, 4000, 10000}), Figure 3.

**Figure 3.**
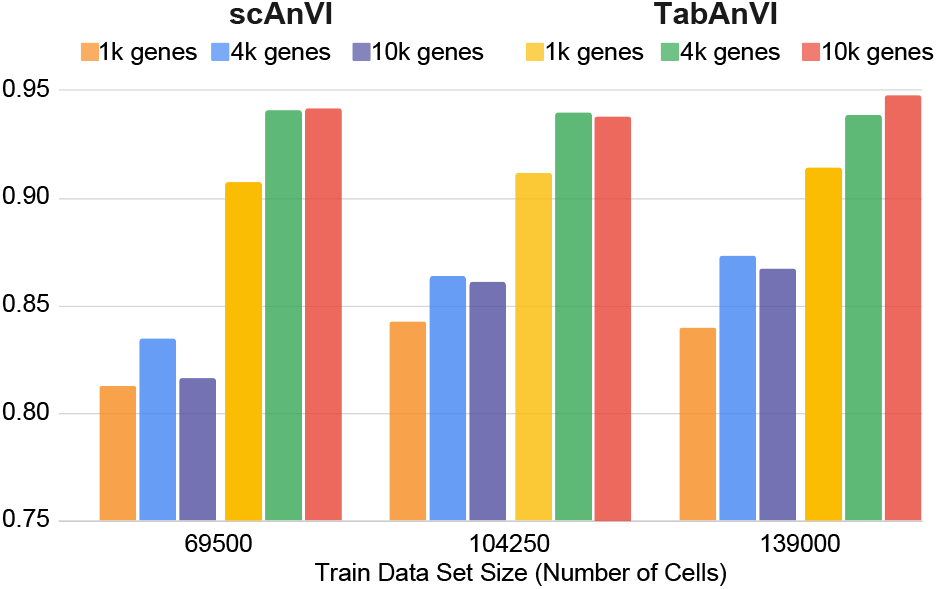
Scalability performance evaluated on subsets of features and observations. We compare macro F1 annotation metrics from TabVI against scAnVI when the training data consists of fewer genes, fewer cells, or both. TabVI performance consistently improves when more features are introduced in the input data. Each data point is repeated 3 times; we display mean values.

TabVI demonstrates strong performance even when the dataset is subsampled. In contrast, scAnVI under-performs in all settings. Moreover, when the number of input features was expanded from 1,000 to 4,000 genes, TabVI’s performance improved significantly across all data settings. For completeness we also ran the model with 10,000 genes, closer to the order of magnitude available in genomics data, and saw no significant improvement over using the top 4,000 genes. We attribute this improvement in performance to the use of sample-specific transformer attention masks. In contrast, scAnVI either experienced a decline in performance or showed no notable improvement with the increased feature set. In the next section, we will explore the output of attention masks to interpreted their values given the input data.

Taken together, these computational experiments demonstrate that the inclusion of a tabular transformer-based architecture in a probabilistic deep generative model lends improvements in the representation of single-cell data sets. Notably, our tabular transformer architecture shows potential as a sample-efficient approach, performing robustly across varying data and feature sizes.

### 4.3 Meaningful Cell Type-specific Feature Selection

To better understand the mechanism by which sample-specific attention masks enhance performance in our tasks, we analyze the attention values produced by the attentive transformer (Figure 1) for the neurons within the Sst subclass, a key population of cortical inhibitory neurons. This subclass is the earliest neuronal population vulnerable to Alzheimer’s Disease pathology Gabitto et al. (2023); E. et al. (2022). The Sst subclass contains 16 unique cell types, each containing 1,000 examples in our studied dataset. To better visualize which genes are attended to, we select a threshold to identify the most significant attention values.

This threshold is defined as the approximate inflection point in the ranked average attention mask values for each gene across all examples in the Sst subclass (Figure 4a). Next, to identify a number of highly attentive genes, we examine the inflection point of the ranked fraction of attentive cells per gene (Figure 4b). This thresholding process identifies 200 genes deemed highly attentive in the Sst subclass.

**Figure 4.**
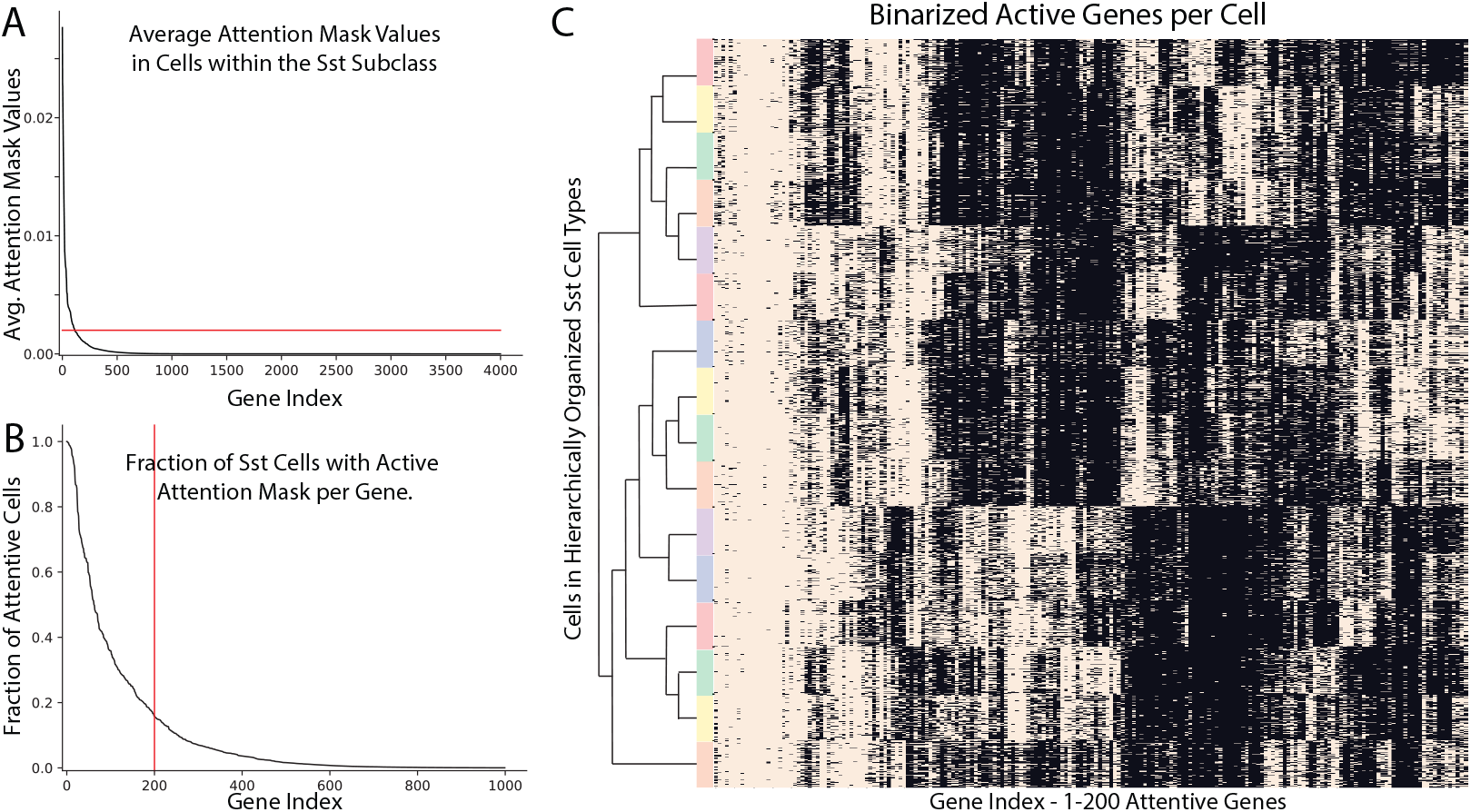
The transformer attention mechanism captures cell type-specific yet combinatorial attention patterns. **(a)** Line plot depicting the value at the output of the attention transformer, average across all Sst cells (y-axis) for all input genes (x-axis). Values are sorted for visualization purposes. The red line marks the threshold for genes to be considered “active” or “being paid attention to”, set at the inflection point of the attention mask values. **(b)** Line plot depicting for each gene (x-axis) the fraction of Sst cells in which the gene is active or being paid attention to (y-axis), interpreted as the probability that a gene is attended to, given the input data. Genes are sorted by decreasing attention probability. **(c)** Heatmap illustrating the attention mask values for all cells within the Sst subclass, arranged hierarchically by genes (x-axis or columns) and stratified by cell types within the Sst subclass (y-axis or rows). With color denoting supertypes within the Sst subclass.

TabVI produces highly selective attention masks at both the subclass and supertype levels (Figure 4c). These mask-attentive features can be divided into three categories: those that are inactive, those *consistently active* at the subclass level (left, Figure 4c), and those that are *cell type-specific* (middle and right, Figure 4c). Cell type-specific masks are active across various cell types, reflecting the granular nature of cell type labels, where no single gene uniquely defines a cell type. A closer analysis of gene expression within these masks allows us to distinguish between masks associated with sparsely expressed genes and those linked to genes broadly expressed across cells. Sparsity is a well-known challenge in single-cell datasets G. et al. (2023). Sparsely expressed genes contribute minimally to cell type characterization, as they may be highly selective but are only present in a small number of cells within each type A.5. In contrast, broadly expressed genes exhibit differential expression across cell types and demonstrate strong selectivity across multiple cell types. In summary, these findings show that granular cell type labels are not defined by a single gene but rather by combinations of genes. Cell type annotation algorithms can enhance their accuracy by learning multiple gene patterns in a hierarchical fashion, which vary across subclasses.

## 5 Discussion

Here we introduced TabVI, a deep generative model for the analysis and annotation of single-cell RNA-seq data that harnesses a lightweight and sample efficient tabular transformer architecture to enhance performance on downstream tasks. Using integration and cell type classification benchmarks, we showed that TabVI improves annotation accuracy compared to baseline probabilistic models (scVI/ScAnVI) and NLP-based transformers (scGPT) while retaining latent space interpretability and batch correction capabilities. Future work will focus on studying generalization capabilities of our tabular transformer architecture across vast data sets and out-of-sample examples while extending our model architecture to multimodal single cell assays.

## A Appendix

### A.1 TabAnVI Probabilistic Model

In the case of TabAnVI, the model is enriched with a discrete cell type label *c*_*n*_, having a multinomial prior, and a hierarchy on the latent biological states *u*_*n*_ and *z*_*n*_. The full probabilistic model for TabAnVI is described in equation 4.

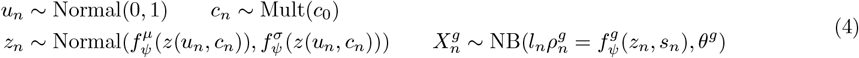

where in the previous equation we have overloaded notation to collect all parameters from all nonlinear functions *f*^*µ*^, *f*^*σ*^, and *f*^*g*^ into *ψ*. These functions act as decoders within the variational autoencoder framework, mapping from *u*_*n*_ to *z*_*n*_ and then to 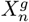. To prevent posterior collapse, a widely studied phenomenon where the posterior of the latent variables is equal to the prior Wang et al. (2023); Bowman et al. (2016); Chen et al. (2017), we limit the decoder capacity in TabAnVI and use fully connected networks (similarly to ScAnVI).

To infer posterior parameters, we performed approximated Bayesian inference within the variational autoencoder framework.

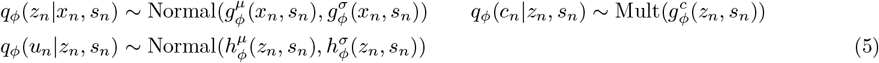

where we have overloaded the notation for all function parameters for *g*^*µ*^, *g*^*σ*^, and *g*^*c*^ into variable *ϕ*. The nonlinear function *g*^*µ*^ act as an encoder within the VAE framework as they map input data *X*^*g*^ into latent variables *z*_*n*_, and are parametrized with fully connected neural networks in scAnVI and tabular-transformers in TabAnVI. In both models, the network that allows cell type classification is parametrized with fully connected neural network architectures.

### A.2 Tabular Transformer Architecture

In the following description, all fully connected layers FC and gated linear units GLU operate per-sample, while batch normalization BN operates over the samples in the batch as usual. The general feature transformer, *f*, maps the input *x* ∈ ℛ ^*B×G*^ to an embedding *Y* ∈ ℛ ^*B×*(*D*+*A*)^. *f* is composed as neural network blocks FC → BN → GLU with residual connections as shown in Figure 1B.

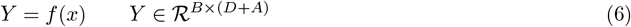

This embedding is split into two parts, *Y*_*D*_ and *Y*_*A*_ (of dimensions *B × D* and *B × A*, respectively). *Y*_*D*_ is used to transform the input, while *Y*_*A*_ is used as input to the attention transformer in computing attention masks. We set *A* and *D* as hyperparameters.

The attention transformer, *g*, maps *Y*_*A*_ ∈ *R*^*B×A*^ to *R*^*B×G*^, creating mask *M*. *g* is composed of blocks FC → BN (Figure 1C).

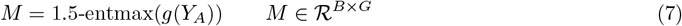

Using 1.5-entmax Peters et al. (2019) ensures sparsity in the learned masks, and that ∑_*g*_ *M*_*bg*_ = 1.

The encoder architecture converts input count values, *X*∈ ℛ^*B×G*^, to normalized and log-transformed values for each sample (cell), *X*_norm_ ∈^*ℛB×G*^. This input is passed through the feature transformer, *f*, which outputs an embedding *Y* ^1^. The embedding is split into 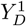 (discarded in this step) and 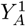, which is passed to the attention transformer.

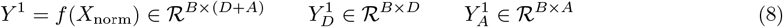

The attention transformer *g* is applied to 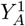, producing the attention mask *M*. This mask is then applied element-wise to the input *X*_norm_, producing the masked input 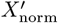.

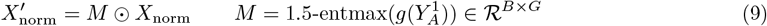

The updated input 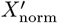 is passed through the feature transformer *f* again, resulting in embedding *Y* ^2^. This is split into 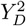, which is used for subsequent processing, and 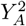, which is discarded as we only consider a single decision step, as in prior methods Fischer et al. (2024).

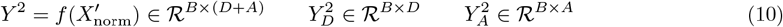

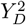 is passed through a fully connected (FC) layer to generate the posterior mean *µ* and variance *σ*^2^ as output of the encoder.

### A.3 Dataset

We consider a human middle temporal gyrus snRNA-seq dataset, which consists of 139k cells originating from 84 donors, spanning the entire spectrum of Alzheimer’s disease states Gabitto et al. (2023). The dataset contains hierarchically organized cellular type annotations, with broad, aggregated *subclasses* and finer, granular *cell types*. There are 27 subclasses and 139 cell types.

Following standard workflows and to reduce computational costs of benchmarking, we use 1,000, 4,000, and 10,000 highly variable genes in training and analysis, calculated based on gene expression count values, treating donors as the batch variable. The train and test sets are computed by selecting 80% and 20% of the examples in each cell type, respectively. When sub-sampling the dataset (4.2), we select 50% and 70% of the cells in each cell type evenly. In subsequent analysis, dataset size refers to the total number of cells in both train and test sets.

### A.4 Model training

scVI and TabVI are first pre-trained for 500 epochs, then model weights are loaded into their corresponding extensions (scAnVI and TabAnVI) and fine-tuned on labeled data for 35 epochs without freezing the latent space. ScVI and TabVI refer to the pre-trained model, while scAnVI and TabAnVI refer to the fine-tuned model. Cell type annotation metrics are computed using the final fine-tuned model as it is the only one with a cell type annotation layer within its infrastructure. Attention masks are computed through one forward pass of the pre-trained TabVI and thus reflect the model’s attention during reconstruction of the original counts data. All models were trained on NVIDIA A100 Tensor Core GPUs.

Model parameters are reported in table 1. Decision steps refer to the number of iterations in which the attentive architecture is repeated. Attention shared layers refers to the number of shared, fully connected layers in the encoder. Attention independent layers refers to the number of independent GLU layers. The decoder is a multi-layer perceptron (MLP) with its own set of FC layers.

**Table 1:** Model Parameters.

| Parameter | TabVI | scVI | TabVI-Attention Only |
| --- | --- | --- | --- |
| Batch Size | 128 | 128 | 128 |
| Learning Rate | 1.00E-04 | 1.00E-04 | 1.00E-04 |
| Optimizer | AdamW | AdamW | AdamW |
| Weight Decay | 1.00E-06 | 1.00E-06 | 1.00E-06 |
| Pretraining Epochs | 500 | 500 | 500 |
| Fine-tuning Epochs | 35 | 35 | 35 |
| $n_a$ | 64 | N/A | 64 |
| $n_d$ | 128 | N/A | 128 |
| Mask Type | 1.5 Entmax | N/A | 1.5 Entmax |
| Decision Steps | 1 | N/A | 1 |
| Attention Shared Layers | 3 | N/A | 3 |
| Attention Independent Layers | 5 | N/A | 5 |
| FT Blocks | 2 | N/A | 1 |
| AT Blocks | 1 | N/A | 1 |
| Activation | GELU | ReLU | GELU |
| FC Layers Decoder | 3 | 3 | 3 |
| Latent Dimension | 20 | 20 | 20 |
| Hidden Dimension | 256 | 256 | 256 |

### A.5 Cell Type-specific Feature Selection

To better understand the selectivity of our attention mechanism, we separate the 200 most active genes into sparsely expressed and non-sparsely expressed sets. Once again, we consider the Sst subclass for analysis. Expression values are first scaled to 10,000 counts per-example, then log1p transformed. To make the distinction between sparse and non-sparse genes, we normalize the expression values per-gene, then sum the expression values per-gene, taking the top 1% of genes as non-sparse and the remaining genes as sparse. This yields 72 non-sparse genes and 128 sparse genes. The expression values in both cases are highly selective of cell type, though this pattern is much more visible in the non-sparse expression map (Figure 5).

**Figure 5.**
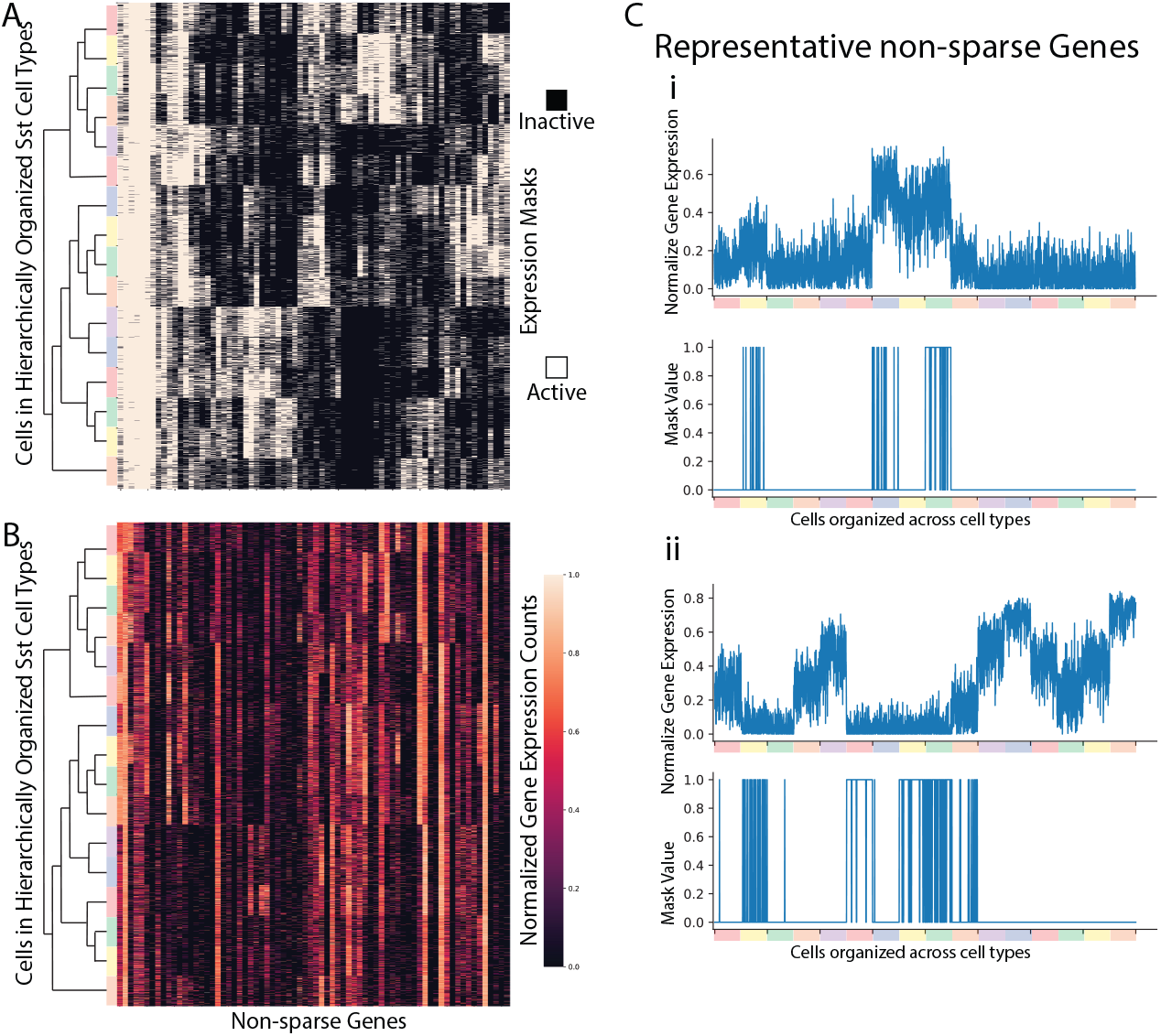
Transformer attention masks select combinatorial, highly selective genes. **(a)** Heatmap illustrating attention masks corresponding to non-sparsely expressed genes for all cells within the Sst subclass, arranged hierarchically by genes (columns) in the mask and hierarchically stratified by cell types within the Sst subclass (rows). **(b)** Heatmap illustrating normalized expression values corresponding to sparsely expressed genes. Cells and genes organized as in **a. (c)** Two representative non-sparse genes with their corresponding masks display as a line plot.

### A.6 Studying Encoder Architecture - Ablation Experiments

To better understand the impact of the FT and AT blocks within the network, we conduct a single ablation experiment. We compare the performance of TabAnVI when using two encoder configurations, the first one termed TabAnVI, uses the entire architecture (section A.1). The second one, termed TabAnVI-Attention only, in which only the attention transformer layer is part of the architecture and not the second feature transformer. Parameters for this model are included in table 1. We compare models trained on 1k, 4k, and 10k genes across 50%, 75%, and 100% of the 139,000 cell SEA-AD dataset. We measure performance using mean F1 score. We show the performance of TabAnVI, TabAnVI AT only, and scAnVI in figure 6.

**Figure 6.**
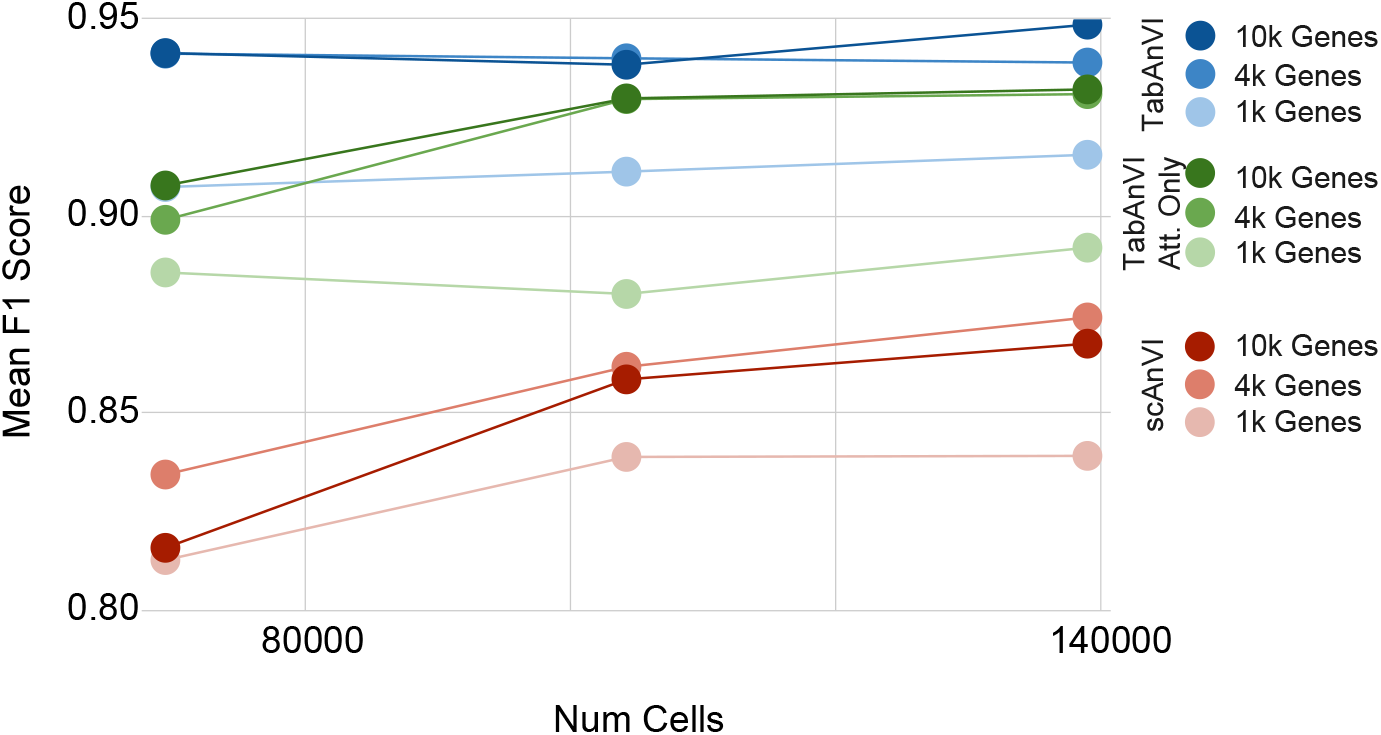
All Components of our attention architecture are necessary to increase annotation performance. Performance comparison of TabAnVI, TabANVI Attention only, and scAnVI across multiple data and feature settings. Models were trained on subsets (50%, 75%, and 100%) of the SEA-AD dataset and 1k, 4k, and 10k genes. The ablation study illustrates the impact of the FT and AT blocks within TabAnVI’s architecture

## Notes

### Competing Interest Statement

The authors have declared no competing interest.

